# Emotional Responses to Naturalistic and AI-generated Affective Pictures: A Systematic Comparison

**DOI:** 10.1101/2025.11.03.686276

**Authors:** Faith Gilbert, Fan Yang, Ethan Smith, Sarah Gardy, Andrew Farkas, Mingzhou Ding, Ruogu Fang, Andreas Keil

## Abstract

Pictures depicting naturalistic scenes are widely used in studies of human emotion. However, the practical use of affective pictures is limited by several factors, including the difficulty of obtaining content-diverse, high-quality, openly accessible, and standardized stimuli that are necessary for specific research questions. The use of artificially generated (AI) pictures could address this limitation, but it is unclear if AI-generated pictures evoke reliable emotional responses. The present study sought to address these challenges by comparing emotional responses to AI-generated pictures with responses to original, standardized, pictures. In two study iterations, standardized pictures containing pleasant, neutral, and unpleasant content were selected from the International Affective Picture System (IAPS) and other sources. Then, a matched AI-counterpart was created for each original picture using generative deep neural networks. A total of 109 participants viewed the picture sets while pupil diameter and electroencephalogram (EEG) were recorded. Evaluative ratings of hedonic valence and emotional arousal were also collected. For both AI and original exemplars, pictures depicting emotional content elicited stronger responses than neutral content for ratings and EEG-derived variables, with weaker effect sizes for the AI-generated pictures. Furthermore, picture-level analyses found that ratings and EEG measures were strongly correlated between matched AI and original pictures. Pupil data also showed the expected content effects in study iteration 2, but not in iteration 1. Together, this initial study suggests that AI-generated picture sets can effectively elicit well-established self-reported affect and physiological responses, presenting a promising avenue for future studies of human emotion.

## Introduction

Viewing affective pictures is a widely used paradigm for inducing and investigating emotional responses in human observers (Lang et al., 1993; Radilová, 1982). Previous studies have shown that such pictures elicit strong autonomic, visceral, and neural responses that vary as a function of picture category, with emotional content (categorized broadly as pleasant and unpleasant) eliciting stronger responses than pictures containing neutral content (Bradley et al., 2008; Codispoti et al., 2001, 2006). The most widely used collection of standardized emotional pictures is the International Affective Picture System (IAPS, Lang et al., 1997) although there are various other sets (e.g., Kurdi et al., 2017; Marchewka et al., 2014).

However, despite the experimental control and consistency offered by such picture sets, they often come with several limitations. First, pictures can quickly become outdated (Marchewka et al., 2014). As cultural and societal norms change, pictures lose their potency or realism, particularly if they depict outdated features such as hairstyles, technology, or fashion. Second, using pictures also poses a challenge in terms of quantity and content specificity: Developing and validating a picture set that is appropriate for studying a particular facet of emotional experience and behavior requires the collection, standardization, and curation of pictures. In this process, researchers typically select exemplars from established databases, capture and edit new pictures, or download pictures from an online source, and obtain copyright. These steps are often followed by quantifying and controlling for low-level physical features and cross-validation against behavioral and physiological measures. Although there are ample pictures in such sets for broader unpleasant or phobic categories, such as gore, snakes, and mutilation (Dan-Glauser & Scherer, 2011), there is a distinct lack of pictures applicable to more specific research problems, including clinically relevant content, such as contamination or body image. Moreover, as many facets of psychophysiological work require extensive amounts of trials, the pictures found in these categories are often not numerous enough to accommodate this need. This necessitates the collection of more specific content from multiple different affective sets, which takes a considerable amount of time and resources (Dan-Glauser & Scherer, 2011; Kurdi et al., 2017).

Because of these limitations, there is a pressing need for an adaptive picture system capable of generating a large amount of controlled and validated content that is suitable for a wide range of applications. A potential solution is the use of pictures generated from artificial intelligence (AI) models. This approach would save not only time but also allow for greater picture specificity due to the flexibility of prompt inputs. However, the use of AI-generated pictures in affective science remains relatively unexplored. The present study seeks to examine the extent to which pictures generated from AI models can evoke well-established behavioral and physiological responses.

Decades of research have shown that pictures evoke reliable emotional responses (Bradley et al., 2022; Codispoti et al., 2001; Lang et al., 1993; Radilová, 1982). Pictures are often grouped into categories of pleasant, neutral, and unpleasant content (e.g., Schupp et al., 2006). Previous examinations of physiological responses to affective pictures have shown greater reactivity to emotional as opposed to neutral content. One such response is the late positive potential (LPP), an event-related component that is larger for pleasant and unpleasant pictures when compared to neutral pictures (Olofsson et al., 2008). This effect has been most commonly observed from approximately 400-800 ms after picture onset in centro-parietal electrodes (Codispoti et al., 2006; Pastor et al., 2008). Likewise, when examining the spectral power of the EEG signal, alpha band power (8-13 Hz) decreases more for emotional relative to neutral pictures (Codispoti et al., 2023). This finding has been interpreted as a marker of saliency-driven increases in sensory processing, consistent with concepts such as selective attention (De Cesarei & Codispoti, 2011; Schubring & Schupp, 2019). Finally, increases in pupil diameter to emotionally relevant pictures are a measure of sympathetic arousal (Bradley et al., 2008; Reimer et al., 2016).

The present study sought to explore the potential of AI-generated pictures by comparing their emotional responses against pictures taken from the IAPS and other validated sources. Here, affective responses were examined using three physiological indices (LPP, alpha power change, pupil diameter) along with self-reported hedonic valence and emotional arousal. Each AI-generated picture was derived from a matched original naturalistic picture. Behavioral and physiological responses to both original pictures and the AI set were then compared. Given the literature above, it was hypothesized that both original and AI-generated pictures would elicit well-established patterns of heightened physiological responses and self-reported affect to emotional, compared to neutral, pictures.

## Material and Methods

### Participants

This study was comprised of two separate iterations, organized in a between-subjects design. All participants were healthy and had no history of seizures or recent concussions. Prior to the experimental session, participants were informed that they would be viewing pictures ranging in emotional content, some of which could be disturbing. However, they were not told about the origin of the pictures (i.e., AI-generated or other). After consenting, participants were fitted with an electroencephalogram (EEG) net and completed the picture viewing task. Following their participation in the study, participants were compensated with course credits. This study was approved by the University of Florida’s institutional review board and was conducted in compliance with the Declaration of Helsinki.

In the first iteration, 45 participants contributed to EEG data, 61 to behavioral rating data, and 59 to pupil dilation data. The 16 participant datasets that were not included in the analyzed sample were due to collection or processing issues including an amplifier malfunction rendering EEG data unusable, participant requests to complete the task without EEG, and a stimulation program malfunction, resulting in incorrect event markers. In total, 61 individuals participated (45 female, 16 male; 45 women, 15 men, 1 non-binary; 38 white, 8 black or African-American, 10 Asian, 3 other, 1 preferred not to answer, 1 of two or more races; 18 Hispanic, 43 non-Hispanic) ranging from ages 18-22 (mean = 19).

In the second iteration, 43 participants contributed to EEG data, 47 to behavioral rating data, and 46 to pupil data. For this iteration, 5 datasets were not analyzed due to collection or processing issues including an amplifier malfunction, rendering EEG data unusable and a stimulation program malfunction, resulting in incorrect event markers. In total, 48 individuals participated (41 female, 7 male; 41 women, 7 men; 36 white, 3 black or African-American, 7 Asian, 1 other, 1 prefer not to answer; 12 Hispanic, 36 non-Hispanic) ranging from ages 18-30 (mean = 19).

### Stimuli

Participants were presented exemplars from IAPS as well as pictures with similar content, curated for prior studies (Farkas et al., 2020). Pleasant pictures depicted content such as victorious athletes, pleasant animals, and romance; neutral pictures included nature scenes, objects, abstract shapes, and neutral animals; while unpleasant pictures included injury and mutilation, threatening humans and animals, and disgust scenes. A rigorous standardization process was used for each AI-picture making an effort to control and standardize other factors such as picture resolution, color composition, and scene composition between matched comparisons. Although the long-term goal of this project is for generative models to produce ready-to-use outputs for real-world applications, the present study focused on the use of AI-pictures as affective stimuli.

In the first iteration, exemplars were selected based on the quality of picture resolution as well as the realism of generated counterparts. Original pictures were utilized as inputs into a stable diffusion model. In this model, text descriptions were generated based on original pictures, which were subsequently input as prompts to produce new pictures. Pictures were generated by inputting keyword descriptions into GPT-4 from which labels (emotion type, label, or other outputs) were generated. These labels were then fed back into GPT-4 to create prompts, after which stabilityAI/stable-diffusion-2-1 was applied to generate a new picture based on the prompt. The resulting pictures were manually filtered and checked for realism based on factors such as prototypical features and naturalistic content. All pictures, original and recreations, were then sized to 512 × 384 pixels and standardized for luminance using functions from the Matlab image processing toolbox, version 2022b. The original pictures category contained 20 pleasant, 20 neutral, and 20 unpleasant pictures, and all originals were matched with corresponding AI counterparts, resulting in a total of 120 pictures.

In the second iteration, exemplars were selected based on the quality of picture resolution as well as normative ratings. Captions for the original IAPS images were first generated using Bootstrapping Language-Image Pre-training (BLIP), a vision-language pretraining framework (BLIP; Li et al., 2022). Based on these captions, hand-designed prompts were produced and fed into a Stable Cascade model to generate high-resolution, realistic pictures (Pernias et al., 2023). All pictures were sized to 900 × 900 pixels and standardized for luminance. The exemplar set contained 40 pleasant, 40 neutral, and 40 unpleasant original pictures respectively, and all original pictures were matched with AI counterparts, resulting in a total of 240 pictures.

### Study design

In both iterations of this study, continuous EEG data as well as metrics of pupil dilation were collected while participants viewed pictures presented on a Display++ LCD Monitor (Cambridge Research Systems Ltd., Rochester, UK) with a screen refresh rate of 120 Hz. All experimental procedures were coded using functions from Psychtoolbox-3 (Kleiner et al., 2022).

In the first iteration of the study, participants were seated at a distance of 120 cm from the screen, with pictures spanning a horizontal visual angle of ∼ 9*°*. Pictures were presented on a black background for 1000 ms followed by a jittered inter-stimulus-interval (ITI) of 2,000-4,000 ms, as shown in Figure 1. Original and AI-generated pictures were presented in a randomized order, where participants would rate each picture directly following its presentation. Trials were split into 2 blocks, with each picture presented twice (once in each block).

**Figure 1.**
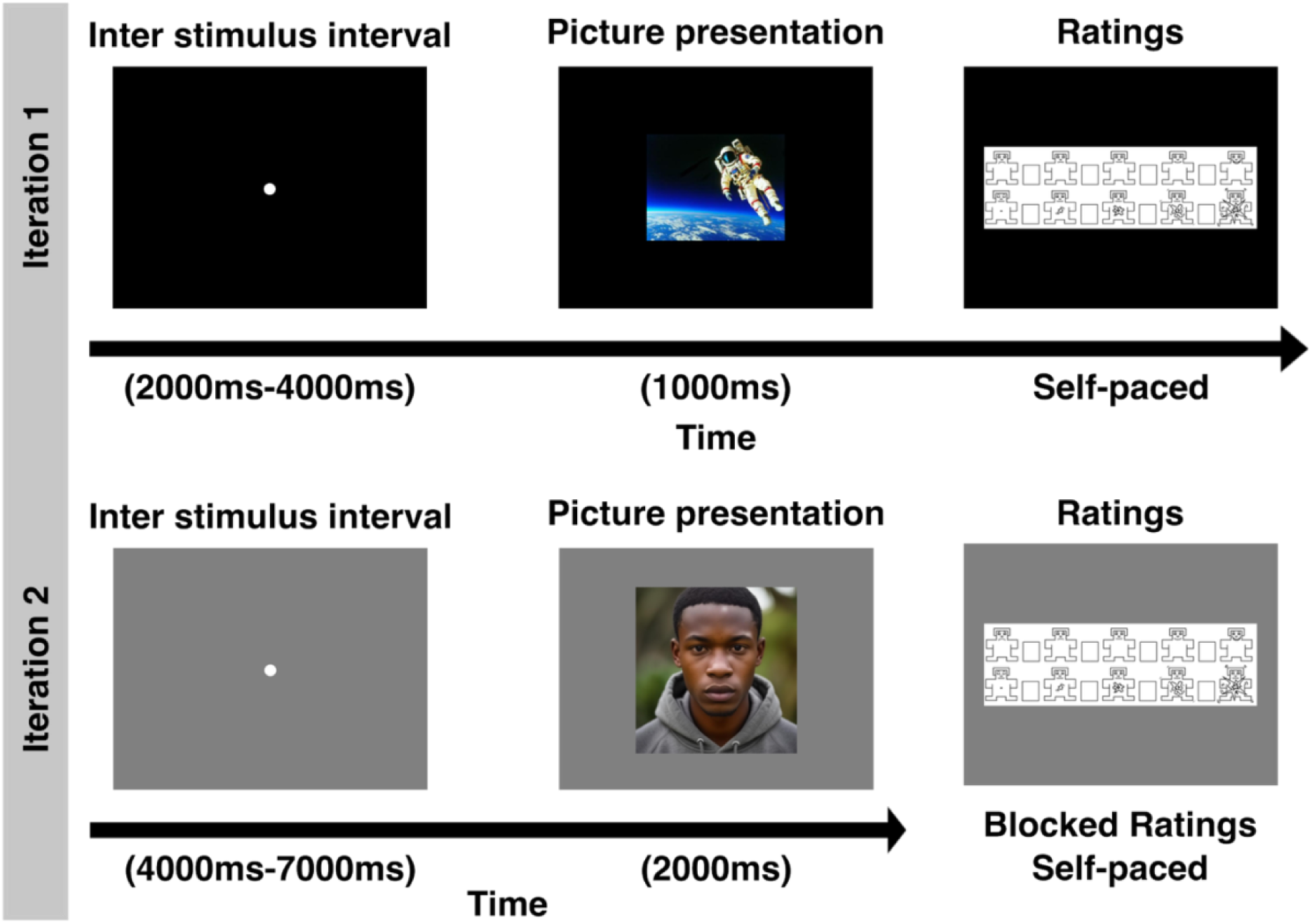
Study design for iterations 1 and 2. In iteration 1, immediately following the picture presentation, participants rated the picture on scales of valence and arousal. The presentation order was randomized for all 120 pictures, and each picture was presented twice. In iteration 2, the presentation order was randomized for all 240 pictures, and each picture was presented once. Following the end of the study, participants rated each picture for valence and arousal in a blocked rating design. Example pictures are taken from the AI-generated stimuli.

In the second iteration, participants were also seated at a distance of 120 cm from the screen, with pictures spanning a visual angle of ∼ 15*°* horizontally and vertically. Initial examination of the first iteration demonstrated a strong light reflex due to the black background, therefore, in the second iteration pictures were presented on a grey background for 2000 ms followed by an ITI of 4000-7000 ms (see Figure 1). Original and AI-generated pictures were presented in a randomized order, with a self-paced break after the first 120 pictures. After all pictures were presented, participants were asked to rate the pictures.

### Ratings

Participants rated all pictures using scales of hedonic valence and emotional arousal from the self-assessment manikin scale. Instructions on how to use the rating scales were given through an adapted script based on the guidelines for the IAPS picture data base (Lang et al., 1997). Ratings of hedonic valence and arousal were then analyzed in two ways: They were averaged within each participant, separately for content categories (pleasant, neutral, unpleasant), and they were also retained for individual pictures for analyses using pictures as observations (see below). In the first iteration, the ratings from the first and second trial were averaged together for both the condition averages, as well as individual pictures. In the second iteration, each picture was only rated once.

### Pupil recording, pre-processing, and visualization

Pupil diameters were measured using an EyeLink 1000 Plus eye tracker system with a 16 mm lens. To calibrate and validate the device, participants were asked to follow a white circle with their gaze, which was presented on a twelve-point grid. Pupil data were sampled at 500 Hz, and the diameter of the pupil was calculated by fitting an ellipse to the mass threshold of the pupil. In preprocessing, data lost due to eyeblinks was accounted for through interpolation. Pupil data were processed from -500 to 4000 ms from the picture onset and filtered using a 7.5-1500 Hz Butterworth filter. Average trial data were then baseline adjusted by subtracting the mean of the baseline (-500 to -2 ms prior to trial onset) from all data points. All data were converted from arbitrary units to mm following calibration.

Pupil diameter was quantified as the mean baseline-adjusted pupil diameter from 1000 to 3000 ms in the first iteration, due to the high contrast between the black background and pictures. As discovered previously, a high contrast between background and stimuli results in a greater initial decrease in pupil diameter (Bradley et al., 2008). Thus, in order to observe pupil diameter differences between stimuli, a larger time window is needed. In the second iteration, pupil diameter was quantified as the mean from 1000-2000 ms. Since the background was grey in the second iteration, the light reflex was less than with a black background, leading to a smaller initial decrease in pupil diameter, and thus required a shorter time window to observe changes in pupil diameter between stimuli.

### EEG recording, pre-processing, and visualization

During the affective picture viewing task, EEG data were recorded using a 128-channel HydroCel Geodesic sensor net and a Net Amp 300 amplifier (Magstim EGI, Oregon, US). Impedances were kept below 60 kOhms, and data was sampled at 500 Hz. Using a processing script that utilized functions of EEGLab (Delorme & Makeig, 2004) in combination with functions from the EMEGS suite of programs (Delorme & Makeig, 2004; Peyk et al., 2011), EEG data was preprocessed offline. First, two Butterworth filters with filter orders of 3 and 9 and cut-off frequencies (3dB points) at 0.2 and 25 Hz, respectively, were successively applied to the continuous data. Then trials were segmented into epochs containing 600 ms pre-picture and 2000 ms post-picture onset. Eyeblinks were corrected using a linear regression correction method based on horizontal and vertical EOG electrodes, located at the outer canthi and above and below both eyes (Schlögl et al., 2007).

Artifact rejection was then implemented following the Statistical Control of Artifacts in Dense Array EEG/MEG Studies (SCADS; Junghöfer et al., 2000). To this end, we first quantified, at the levels of channels and trials, the median absolute voltage, the standard deviation, and the maximum derivative of voltage values. These were standardized and linearly combined into a compound data quality (QC) index. In an initial step applied to data from all trials, we then identified channels in which the QC index was above the median of the distribution by more than 2.5 standard deviations. Those channels were then interpolated using spherical spline functions (Junghöfer et al., 2000). The mean number of interpolated channels in the first iteration was 9, with a range of 0-15. In the second iteration, the mean was 8 with a range of 1-16. Then, this step was repeated for each trial separately with the same threshold, resulting in interpolation of channels that were outlying in a given trial. The mean number of interpolated channels for the first iteration was 5 with a range of 0-16, and in the second iteration, the mean was 5 with a range of 0-17. Finally, the compound index for the remaining trials was calculated across channels, and trials that were 1.25 standard deviations above the mean of the trial-based QC index distribution were discarded from the analysis. An average of 31 trials per condition were retained for the first iteration (pleasant original = 81% with a range of 47-100%, neutral original = 81% with a range of 58-98%, unpleasant original = 81% with a range of 38-100%, pleasant AI = 80% with a range of 53-100%, neutral AI = 81% with a range of 50-100%, unpleasant AI = 78% with a range of 48-98%), and an average of 33 trials per condition were retained in the second iteration (pleasant original = 82% with a range of 55-98%, neutral original = 80% with a range of 50-100%, unpleasant original = 82% with a range of 58-100%, pleasant AI = 83% with a range of 50-100%, neutral AI = 83% with a range of 55-100%, unpleasant AI = 83% with a range of 55-100%).

EEG data were analyzed through quantifying the amplitude of the LPP component and through quantifying changes in alpha power reduction upon the presentation of the pictures. The late positive potential was quantified as an averaged time window of 400-800ms from a centro-parietal electrode cluster centered on CPz. This was completed for both the condition averages (pleasant, neutral, and unpleasant for AI and original pictures), as well as for individual pictures (see below). Alpha reduction was calculated through convolving the data with a Morlet wavelet with a Morlet parameter (*m*) of 10. At the alpha frequency chosen for measurement (10.3 to 12.6 Hz), the frequency uncertainty sigma(f) was 1.03 Hz and the uncertainty in the time domain sigma(t) was 155 ms. Frequency bins were examined from 3.46 to 30.02 Hz in steps of 1.14, with an intrinsic frequency resolution of 0.38 Hz. Average trial data were then baseline corrected by dividing the time-frequency matrix by the average power measured between -200 to -400 ms prior to picture onset. Alpha decrease was then quantified by averaging the time-varying, baseline adjusted power at 10.3-12.6 Hz across a parieto-occipital cluster containing Pz and its 14 nearest neighbors, and across the time points from 500 to 1500 ms. These electrode sites and time points were determined based on a collapsed localizer, averaging across conditions and participants (Luck & Kappenman, 2013).

### Statistical analysis

#### ANOVAs and linear contrasts

Following a standard statistical approach that has been widely used in this field of research, measures were evaluated using mixed model ANOVA with within-subject factors of type (original or AI) and condition (pleasant, neutral, or unpleasant), and a between-subjects factor of iteration (first iteration vs. second iteration) in JASP (Jasp, 2018). Sphericity controls were applied using a Greenhouse-Geisser correction. As the focus of the current study was to assess the validity of AI pictures as affective stimuli (and not to assess their validity as replicates of original pictures), the present study focused on main effects and interactions involving the factor condition.

To examine the presence of the expected content-related effects in each type and iteration, effect sizes of the expected condition differences were estimated using a contrast analysis (Rosenthal & Rosnow, 1985). The weights used in these analyses were linear (-1, 0, 1) for valence ratings, and quadratic (1, -2, 1) for arousal ratings, pupil, LPP, and alpha power data. Effect size correlations (r) were used to quantify effect sizes for the linear and quadratic trends for each dependent variables, as appropriate, and are reported along with F- and p-values for each planned contrast.

#### Picture-level analysis

Additionally, a picture-level analysis was performed by aggregating ratings, LPP, and pupil data picture-wise across participants, resulting in data vectors containing mean responses by picture. All data were scored in the same manner as the ANOVAs and contrasts, with LPP data scored from 400-800 ms from a centro-parietal electrode cluster centered on CPz, alpha data scored from 500-1500 ms on a parieto-occipital cluster containing Pz and its 14 nearest neighbors, and pupil data scored from 1000-3000 ms in the first iteration and 1000-2000 ms in the second iteration. Then, correlation coefficients were calculated between the vectors containing corresponding responses by picture type (original or AI), resulting in a metric of correspondence between original and AI-generated pictures. This method was used as a supplementary visualization of the correspondence between AI and original pictures. Thus, no p-values are reported for these correlation coefficients, reflecting the fact that they are intended as continuous metrics of correspondence/replicability between AI and original pictures, but are not intended to be used for inference statistics.

### Transparency and Openness

This study was not pre-registered. Source code for the task, AI picture materials, and processed data is available at the corresponding OSF project site: https://osf.io/qrx8u/?view_only=7d397b9346ca44c58395d864864e1bcf. However, original IAPS pictures are not available for distribution due to copyright. This study was supported by funding from the National Institutes of Health (Grant: R01MH125615) as well as from the National Science Foundation (Grant: 2318984). We report how we determined our sample size, all data exclusions (if any), all manipulations, and all measures in the study.

## Results

### Ratings

#### ANOVAs and contrasts

When examining ratings of hedonic valence (displeasure) there was a main effect of type, F(1,106) = 56.3, p < 0.001, and condition, F(2,212) = 811.5, p < 0.001. A contrast analysis of the condition differences for each type and each iteration (see Figure 2) demonstrated that the data fit a linear trend, with the least displeasure observed for pictures containing pleasant content, followed by neutral, and unpleasant content prompting the greatest displeasure reports. This pattern was present in original and AI pictures and in each iteration: First iteration, original, F(1,122) = 766.5, p < 0.001, r = 0.96, and AI, F(1,122) = 792.1, p < 0.001, r = 0.96; Second iteration, original, F(1,94) = 731.9, p < 0.001, r = 0.97, and AI, F(1,94) = 562.4, p < 0.001, r = 0.96.

**Figure 2.**
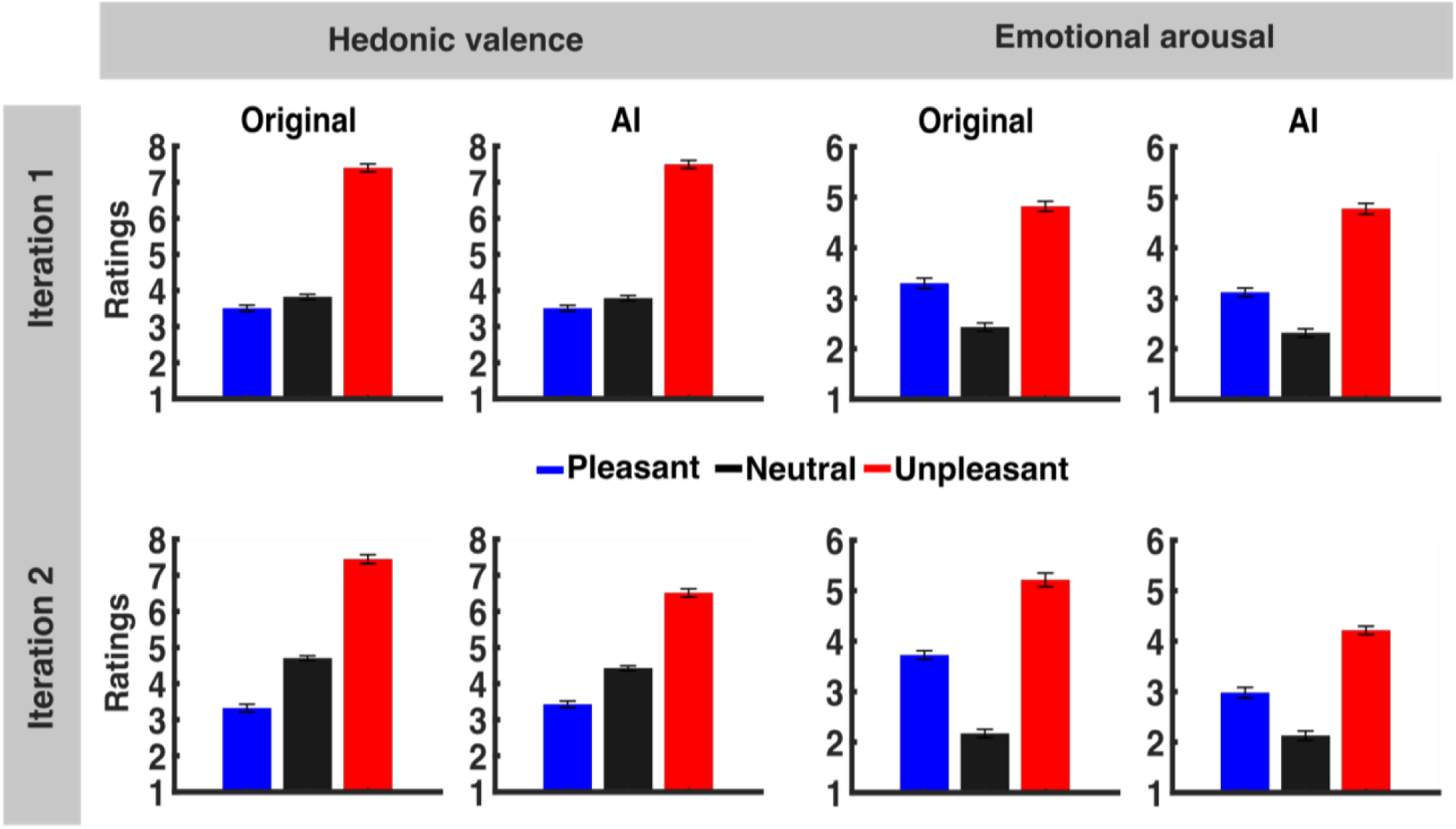
Mean ratings of hedonic valence and emotional arousal for each picture type, condition, and iteration. In the first iteration, each picture was rated twice, while in the second iteration each picture was rated once. The error bars reflect the within-subjects error for each mean.

For arousal ratings, there was likewise a main effect of stimulus type, F(1,106) = 171.3, p < 0.001, and condition, F(2,212) = 295.1, p < 0.001. Arousal ratings followed a quadratic trend, showing a heightened effect for emotional content, where pleasant and unpleasant content were high in arousal, and neutral content was lower in arousal for both original and AI pictures: First iteration, original F(1,122) = 160.2, p < 0.001, r = 0.85, and AI, F(1,122) = 191.5, p < 0.001, r = 0.87; Second iteration, original, F(1,94) = 253.6, p < 0.001, r = 0.92, and AI, F(1,94) = 145.3, p < 0.001, r = 0.87.

#### Picture-level analysis

A comparison between the valence ratings for the different types of stimuli (AI or original) using Pearson’s correlation coefficients revealed a strong positive correlation between the two, as shown in Supplemental Figure 1, for both the first iteration, r = 0.88, as well as the second iteration, r = 0.88. As with valence, there was a strong positive correlation between arousal ratings for AI and original pictures for the first iteration, r = 0.85, as well as the second iteration, r = 0.82.

### Pupil diameter

#### ANOVAs and contrasts

Pupil diameter, as shown in Figure 3, was found to have a main effect of type, F(1,103) = 14.5, p < 0.001, condition, F(2,206) = 37.8, p < 0.001, and iteration, F(1,103) = 302.1, p < 0.001. In the first iteration, there was a fit of a quadratic trend for the original-picture pupil data, but not for the AI-picture pupil data. In the second iteration, the pupil data from the two types (original and AI) had more comparable trends, with both the original and AI demonstrating a strong fit with the expected quadratic trend: First iteration, original, F(1,118) = 25.9, p < 0.001, r = 0.55, and AI, F(1,118) = 0.1, p = 1.713, r = 0.03: Second iteration, original, F(1,92) = 61.8, p < 0.001, r = 0.76, and AI, F(1,92) = 24.8, p < 0.001, r = 0.59.

**Figure 3.**
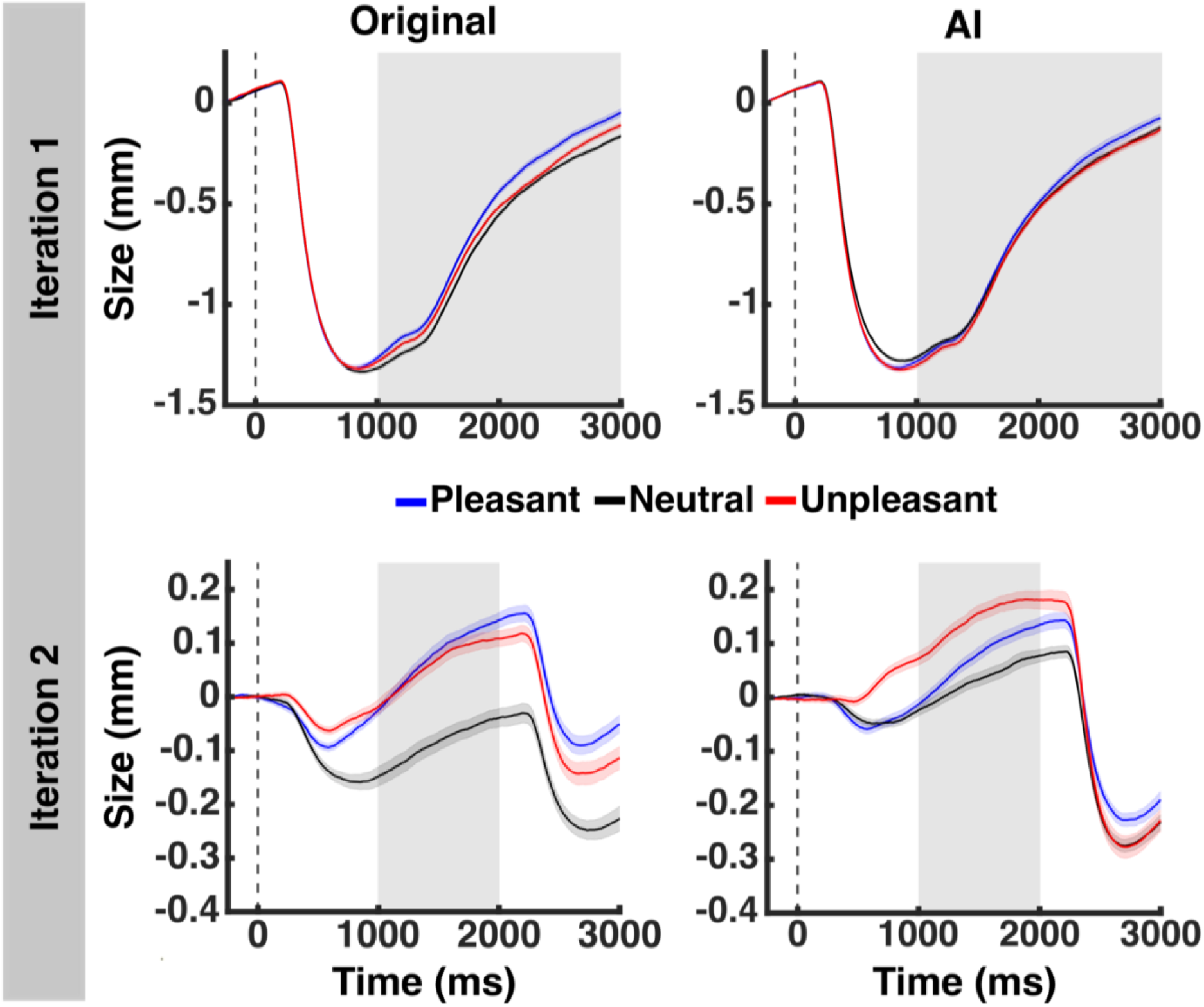
Mean pupil diameter change from baseline over time for each picture type, condition, and iteration. Shaded error bars reflect the within-subjects error for each mean. Different regions of interest were extracted for each iteration, with 1000ms-3000ms extracted for the first iteration, and 1000ms-2000ms extracted for the second iteration, as detailed in the methods section.

#### Picture-level analysis

Paralleling the mean comparisons above, the pupil diameter for the AI and original pictures showed a weak positive correlation for the first iteration, r = 0.27, and a stronger relation for the second iteration, r = 0.44 (see Supplemental Figure 1).

### Late positive potential

#### ANOVAs and contrasts

The LPP amplitude differed between conditions, F(2,172) = 60.1, p < 0.001, and iterations, F(1,86) = 6.2, p = 0.015. A contrast analysis demonstrated that LPP data followed the expected quadratic trend, showing a heightened effect for emotional content (see Figure 4): First iteration, original, F(1,90) = 56.0, p < 0.001, r = 0.74, and AI, F(1,90) = 16.3, p < 0.001, r = 0.52; Second iteration, original, F(1,86) = 71.5, p < 0.001, r = 0.79, and AI, F(1,86) = 27.9, p < 0.001, r = 0.63.

**Figure 4.**
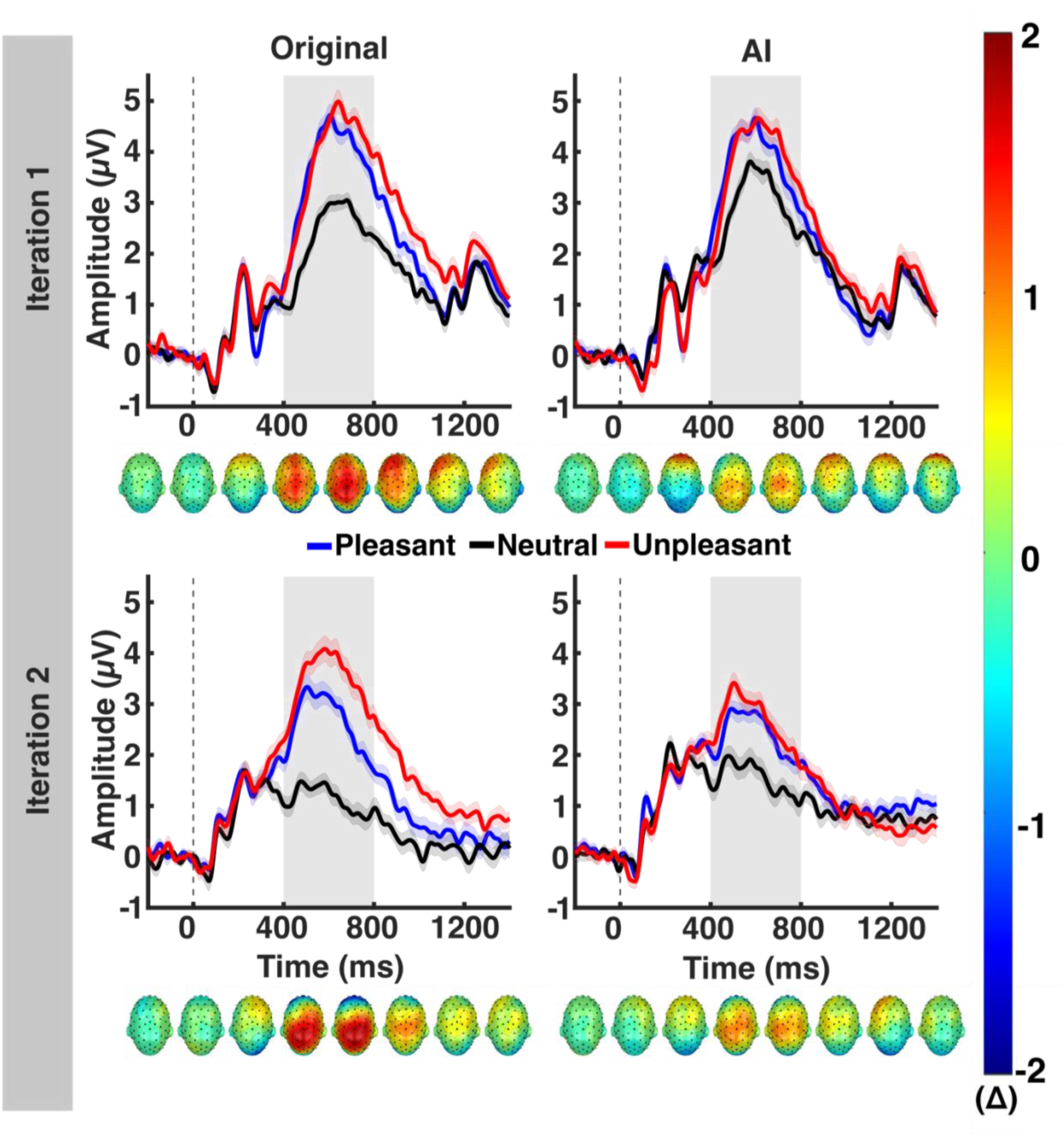
Mean amplitude over time for each picture type, condition, and iteration. Shaded error bars reflect the within-subjects error for each mean. Scalp topographies reflect the difference in amplitude between averaged emotional content (pleasant and unpleasant) and neutral content across time for an averaged cluster of sensors including CPz and its 9 nearest neighbors. The region of interest extracted for analysis is from 400ms-800ms.

#### Picture-level analysis

When examining the relationship between LPP amplitude across picture type, there was a moderate positive correlation between responses to original and AI pictures in the first iteration, r = 0.54, as well as the second iteration, r = 0.56 (see Supplemental Figure 1).

### Alpha power changes

#### ANOVAs and contrasts

Alpha band activity was concentrated from 10-12 Hz around an occipitoparietal cluster, from approximately 500-1500ms, as seen in averaged data in Figure 5. Alpha power differed in response to picture type, F(1, 86) = 5.9, p = 0.017, and condition, F(2,172) = 20.1, p < 0.001 (see Figure 6). In both iterations, there was a strong fit between the data and the quadratic model for the original pictures as well as the AI pictures: First iteration, original, F(1, 90) = 24.8, p < 0.001, r = 0.60, and AI, F(1,90) = 10.6, p < 0.001, r = 0.44; Second iteration, original, F(1,86) = 8.6, p = 0.002, r = 0.41, and AI, F(1,86) = 10.8, p < 0.001, r = 0.45.

**Figure 5.**
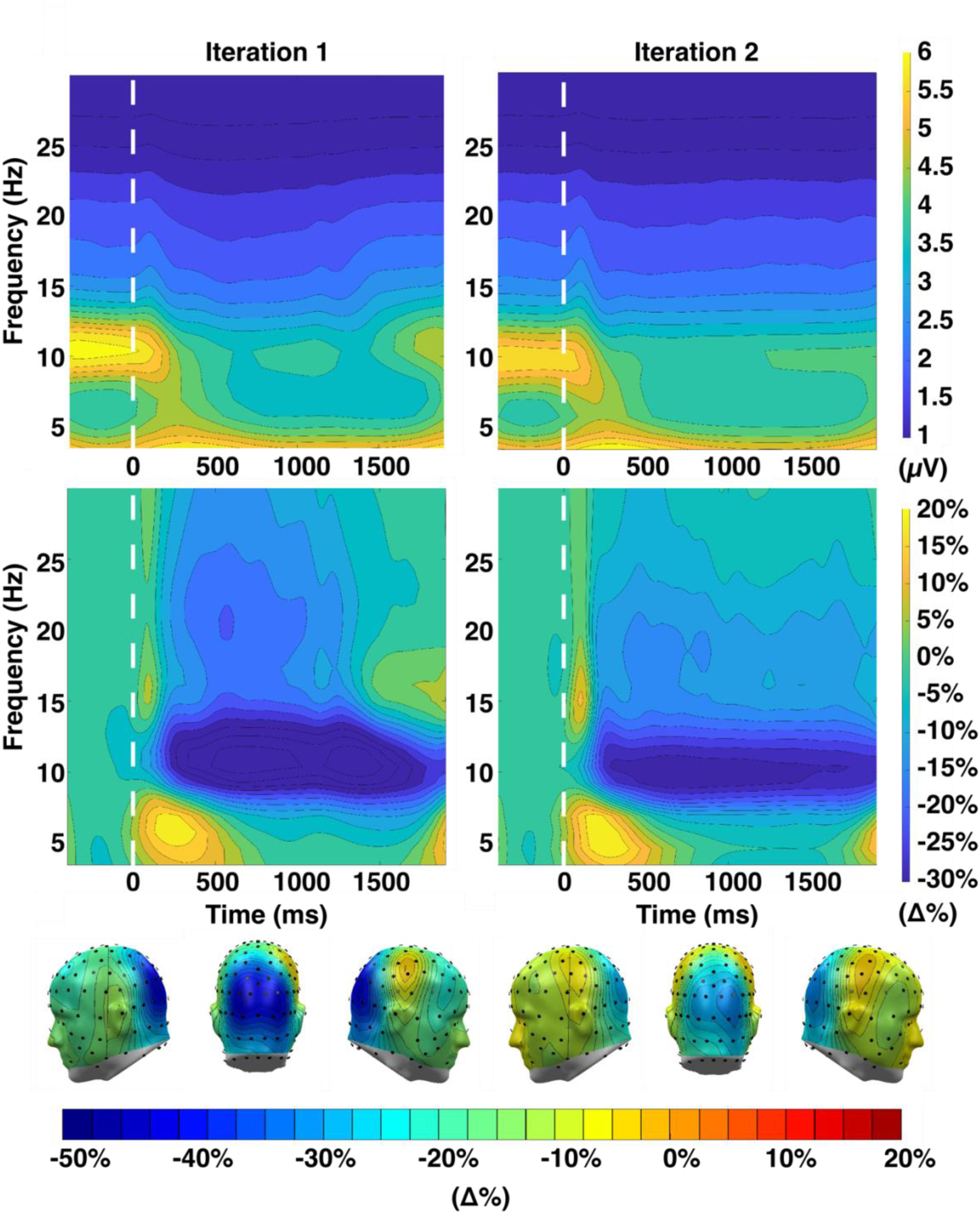
Grand mean alpha band activity for all conditions across time. The top panel represents grand mean power without baseline correction for iteration 1 (N=45) and 2 (N=43), averaged across an occipitoparietal cluster containing Pz and its 14 nearest neighbors. The second panel represents the percentage of power relative to baseline, averaged across sensor Pz and its 14 nearest neighbors. The third panel represents alpha power averaged across 500ms-1500ms and 10-12 Hz.

**Figure 6.**
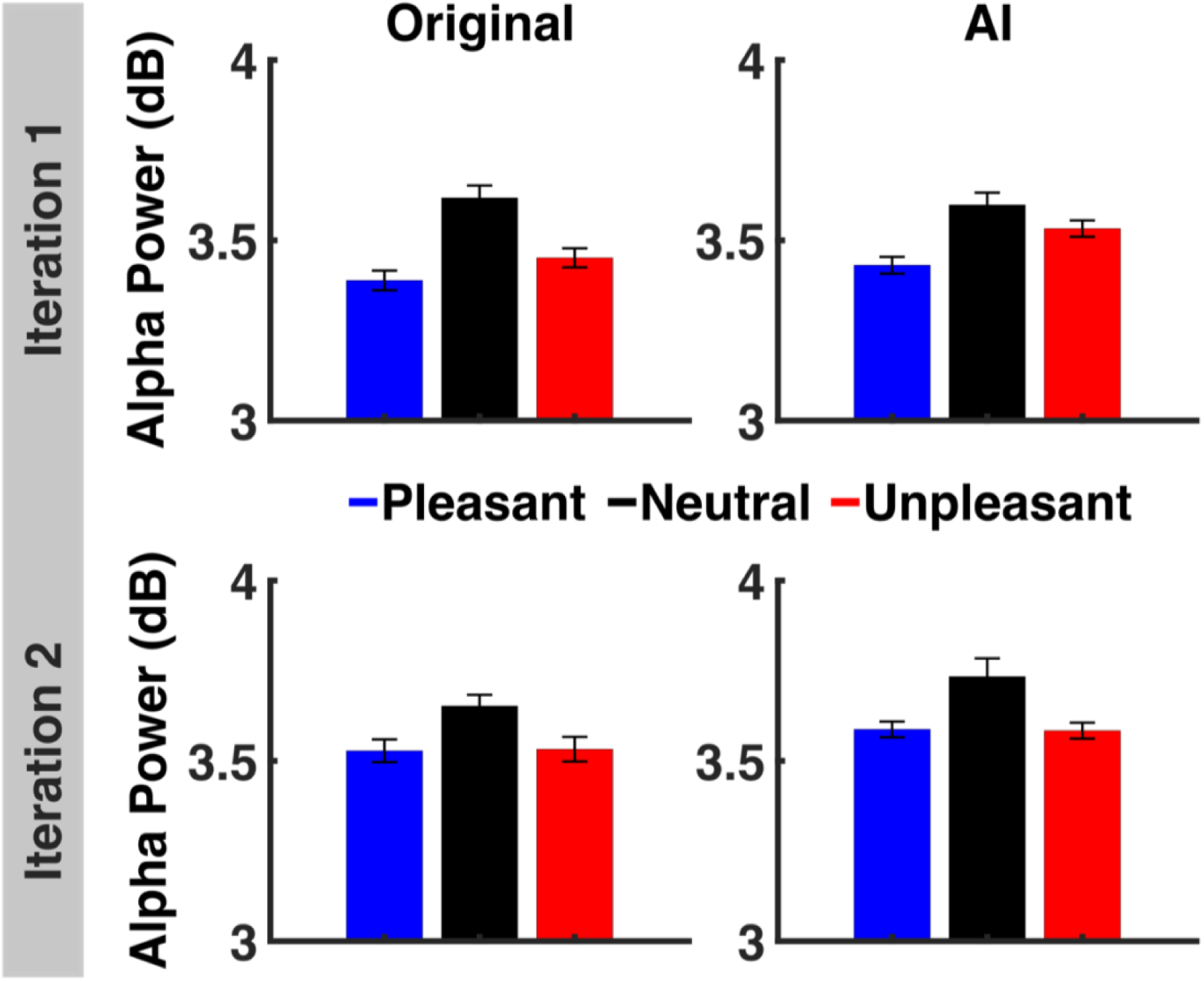
Alpha power in response to pleasant, neutral, and unpleasant conditions for AI and original pictures for each iteration. Alpha power was calculated by averaging across an occipitoparietal cluster containing sensor Pz and its 14 nearest neighbors from 500ms-1500ms and 10-12 Hz.

#### Picture-level analysis

For alpha, there was a moderate correlation between AI and original pictures in the first iteration, r = 0.42, and a weaker correlation between AI and original pictures in the second iteration, r = 0.26 (see Supplemental Figure 1).

## Discussion

A large body of research has demonstrated that viewing emotional pictures elicits stronger physiological and behavioral responses than neutral pictures (Bradley et al., 2001; Codispoti et al., 2001, 2006). However, the usage of existing picture sets comes with societal, cultural, clinical, and technological challenges (Kurdi et al., 2017; Marchewka et al., 2014). The present study sought to address some of these challenges by exploring the reliability and validity of an AI-generated picture set in relation to pre-existing affective pictures. If feasible, a picture set based on an AI platform would allow researchers to generate larger numbers of stimuli, with specific content, on demand, overcoming many limitations of existing sets. In the future, such a platform will likely include algorithms for predicting ratings of hedonic valence and emotional arousal, as initial guidance for users.

In the present study, AI-generated pictures were compared to matched exemplars from well-established picture set to understand if they prompt similar emotional responses to those reported in the existing literature (Bradley et al., 2022). As found in prior studies, there was a systematically greater response to emotional, compared to neutral content. Responses to AI-generated pictures for several dependent variables examined here mirrored responses to their original counterparts. Other variables (pupil diameter, alpha power change) showed some differences between original and AI pictures as discussed below. Overall, however, the present findings suggest that an AI-generated affective picture set may prompt similar response patterns to those observed with existing, carefully curated, affective picture sets, and that many of the widely used responses for assessing human affect do not vary drastically between AI and original pictures.

Comparing different response variables, rating data showed the well-established patterns both for emotional arousal and hedonic valence (Dan-Glauser & Scherer, 2011; Lang et al., 1993) for both picture types (original or AI-generated), which suggests that AI-generated pictures are evaluated similarly to well-curated natural pictures.

In contrast to ratings, pupil data showed more variability in response patterns between iterations and types. In the first iteration, for original pictures, pupil data corresponded with previous findings such that emotional content prompted an increase in pupil diameter compared to neutral content (Bradley et al., 2008). However, this was not the case for AI-generated pictures. The reasons for smaller and more variable effects in the first iteration are manifold and include the smaller visual angle spanned by the pictures, the shorter stimulus duration compared to the second iteration, and the black background resulting in a pronounced light reflex (see Figure 3, top). Together, these factors may have further diminished the effects of AI-generated pictures on the pupil diameter. In the second iteration, effects of AI-generated content were observed in the expected directions, suggesting that with experimental parameters favorable for measuring pupil dilation (gray background, longer presentation times, larger picture size), pupil effects for AI content may more closely mirror effects for the original pictures.

The LPP is a widely used electrophysiological variable in picture viewing studies (Olofsson et al., 2008; Schupp et al., 2006). The current study replicated the typical findings of heightened centroparietal LPP amplitude when viewing emotional content, which we observed for original as well as AI-generated replicates. These results were consistent between iterations. It is noteworthy that differences in LPP amplitude were present for neutral content. Specifically, AI-generated pictures elicited a higher amplitude for neutral content as compared to original pictures. This was observed in both iterations, which suggests that AI-generated neutral content may be more engaging or salient as compared to original neutral pictures. Future studies would benefit from a more detailed exploration as to why AI-generated neutral pictures prompt increased LPP amplitudes as compared to original neutral pictures. An investigation into these differences may also involve a systematic comparison of the semantics contained in on-demand AI-generated neutral scenes versus original neutral scenes.

Finally, alpha power changes differ between emotional and neutral pictures, such that there is a greater decrease in alpha power for emotional pictures compared to the decrease evoked by neutral pictures (Codispoti et al., 2023). In the present study, this pattern was found as well, for the original and AI affective pictures and in both the first iteration and second iteration.

### Future directions and considerations

Across response variables, we largely found correspondence between AI-generated and original pictures, with the strongest correspondence between valence ratings, arousal ratings, LPP data, and alpha data, while the weakest correspondence between original and AI-generated pictures was found for pupil responses in the first iteration. Future studies may systematically explore more expansive AI-generated picture sets, such as those generated without matching exemplars to original counterparts. Additionally, a systematic investigation of AI-picture realism may assess observers’ perceived differences between AI-generated pictures and naturalistic pictures, which may help to explain some of the differences observed in the present study.

The present findings highlight the potential of AI-generated pictures for the exploration of new research questions. For example, targeted and gradual changes could be made in the process of picture generation, generating pictures that represent gradients along a dimension of interest, such as systematically increasing contamination, social threat, or mutilation. Such types of systematically varied and controlled picture sets would facilitate studies of physiological and behavioral responses to clinically relevant content. Furthermore, the ability to alter one aspect of a picture across an entire set opens the door for quantifying how specific scene elements impact behavioral or physiological responses.

However, as the use of AI-generated stimuli is relatively new, it is important for researchers to also consider the impacts of the use of such datasets. Models trained on widely collected pictures may produce replicates generated from creative property or that utilize features unique to an individual. Researchers should consider the ethicality of utilizing a model trained on unknowing or unwilling source material, as well as the constraints that should be placed on picture distribution regarding AI-generated picture sets.

### Constraints on generality

The results of this study are limited in generality by the characteristics of the sample, as well as the context of the study. It has been previously documented that participants tend to display differences in response to affective stimuli with age, such that as age increases responses to pleasant stimuli decrease, while responses to aversive stimuli increase (Keil & Freund, 2009). Consequently, as the sample for this study consists largely of undergraduate students, it is likely that in older demographics results may vary from those of the present study. Additionally, as awareness of AI-generated images increases in society, participants may become more sensitive to the quality or realism of AI pictures. This could have unknown impacts on participant responses to AI-affective pictures, which may impact the generalizability of the study with time.

### Conclusion

When examining physiological and behavioral responses to affective pictures, AI-generated replicates produce comparable responses to original pictures, especially for previously established data patterns found in ratings of hedonic valence and emotional arousal, alpha power changes, and LPP amplitude. These findings suggest the potential implementation of an AI-generated affective picture set, with considerations for the impact and ethicality of the use of AI-generated images in research. In general, particularly in cases where access to previous picture sets is limited, or a greater specificity and volume of pictures is needed for the study, the utilization of an AI-generated affective picture set may be a promising avenue for affective science studies.

## Acknowledgements

The authors would like to acknowledge Ricky Cheng and Angie Cordova for their assistance with this project.

## References

Bradley, M. M., Codispoti, M., Cuthbert, B. N., & Lang, P. J. (2001). Emotion and motivation I: Defensive and appetitive reactions in picture processing. Emotion, 1(3), 276–298. 10.1037/1528-3542.1.3.276

Bradley, M. M., Miccoli, L., Escrig, M. A., & Lang, P. J. (2008). The pupil as a measure of emotional arousal and autonomic activation. Psychophysiology, 45(4), 602–607. 10.1111/j.1469-8986.2008.00654.x

Bradley, M. M., Sambuco, N., & Lang, P. J. (2022). Affective Perception: The Power Is in the Picture. In B. Ionescu, W. A. Bainbridge, & N. Murray (Eds.), Human Perception of Visual Information: Psychological and Computational Perspectives (pp. 59–83). Springer International Publishing. 10.1007/978-3-030-81465-6_3

Codispoti, M., Bradley, M. M., & Lang, P. J. (2001). Affective reactions to briefly presented pictures. Psychophysiology, 38(3), 474–478. 10.1111/1469-8986.3830474

Codispoti, M., De Cesarei, A., & Ferrari, V. (2023). Alpha-band oscillations and emotion: A review of studies on picture perception. Psychophysiology, 60(12), e14438. 10.1111/psyp.14438

Codispoti, M., Ferrari, V., De Cesarei, A., & Cardinale, R. (2006). Implicit and explicit categorization of natural scenes. In S. Anders, G. Ende, M. Junghofer, J. Kissler, & D. Wildgruber (Eds.), Progress in Brain Research (Vol. 156, pp. 53–65). Elsevier. 10.1016/S0079-6123(06)56003-0

Dan-Glauser, E. S., & Scherer, K. R. (2011). The Geneva affective picture database (GAPED): A new 730-picture database focusing on valence and normative significance. Behavior Research Methods, 43(2), 468–477. 10.3758/s13428-011-0064-1

De Cesarei, A., & Codispoti, M. (2011). Affective modulation of the LPP and α-ERD during picture viewing. Psychophysiology, 48(10), 1397–1404. 10.1111/j.1469-8986.2011.01204.x

Delorme, A., & Makeig, S. (2004). EEGLAB: An open source toolbox for analysis of single-trial EEG dynamics including independent component analysis. Journal of Neuroscience Methods, 134(1), 9–21. 10.1016/j.jneumeth.2003.10.009

Farkas, A. H., Oliver, K. I., & Sabatinelli, D. (2020). Emotional and feature-based modulation of the early posterior negativity. Psychophysiology, 57(2), e13484. 10.1111/psyp.13484

Jasp, T. (2018). JASP (Version 0.9)[Computer software]. *URL:* Https://Jasp-Stats.Org.

Junghöfer, M., Elbert, T., Tucker, D. M., & Rockstroh, B. (2000). Statistical control of artifacts in dense array EEG/MEG studies. Psychophysiology, 37(4), 523–532. 10.1111/1469-8986.3740523

Keil, A., & Freund, A. M. (2009). Changes in the sensitivity to appetitive and aversive arousal across adulthood. Psychology and Aging, 24(3), 668–680. 10.1037/a0016969

Kleiner, M., Brainard, D., Pelli, D., Ingling, A., Murray, R., & Broussard, C. (2022). What’s new in Psychtoolbox-3? 2007. Perception, ECVP.

Kurdi, B., Lozano, S., & Banaji, M. R. (2017). Introducing the Open Affective Standardized Image Set (OASIS). Behavior Research Methods, 49(2), 457–470. 10.3758/s13428-016-0715-3

Lang, P. J., Bradley, M. M., & Cuthbert, B. N. (1997). International affective picture system (IAPS): Technical manual and affective ratings. NIMH Center for the Study of Emotion and Attention, 1(39–58), 3.

Lang, P. J., Greenwald, M. K., Bradley, M. M., & Hamm, A. O. (1993). Looking at pictures: Affective, facial, visceral, and behavioral reactions. Psychophysiology, 30(3), 261–273. 10.1111/j.1469-8986.1993.tb03352.x

Li, J., Li, D., Xiong, C., & Hoi, S. (2022). BLIP: Bootstrapping Language-Image Pre-training for Unified Vision-Language Understanding and Generation. Proceedings of the 39^th^ International Conference on Machine Learning, 12888–12900. https://proceedings.mlr.press/v162/li22n.html

Luck, S. J., & Kappenman, E. S. (2013). The Oxford handbook of event-related potential components. Oxford university press.

Marchewka, A., Żurawski, Ł., Jednoróg, K., & Grabowska, A. (2014). The Nencki Affective Picture System (NAPS): Introduction to a novel, standardized, wide-range, high-quality, realistic picture database. Behavior Research Methods, 46(2), 596–610. 10.3758/s13428-013-0379-1

Olofsson, J. K., Nordin, S., Sequeira, H., & Polich, J. (2008). Affective picture processing: An integrative review of ERP findings. Biological Psychology, 77(3), 247–265. 10.1016/j.biopsycho.2007.11.006

Pastor, M. C., Bradley, M. M., Löw, A., Versace, F., Moltó, J., & Lang, P. J. (2008). Affective picture perception: Emotion, context, and the late positive potential. Brain Research, 1189, 145–151. 10.1016/j.brainres.2007.10.072

Pernias, P., Rampas, D., Richter, M. L., Pal, C. J., & Aubreville, M. (2023). *Wuerstchen: An Efficient Architecture for Large-Scale Text-to-Image Diffusion Models* (No. arXiv:2306.00637). arXiv. 10.48550/arXiv.2306.00637

Peyk, P., De Cesarei, A., & Junghöfer, M. (2011). ElectroMagnetoEncephalography Software: Overview and Integration with Other EEG/MEG Toolboxes. Computational Intelligence and Neuroscience, 2011(1), 861705. 10.1155/2011/861705

Radilová, J. (1982). The late positive component of visual evoked response sensitive to emotional factors. *Activitas Nervosa Superior*, Suppl 3(Pt 2), 334–337.

Reimer, J., McGinley, M. J., Liu, Y., Rodenkirch, C., Wang, Q., McCormick, D. A., & Tolias, A. S. (2016). Pupil fluctuations track rapid changes in adrenergic and cholinergic activity in cortex. Nature Communications, 7(1), 13289. 10.1038/ncomms13289

Rosenthal, R., & Rosnow, R. L. (1985). Contrast analysis: Focused comparisons in the analysis of variance. Cambridge University Press. https://cir.nii.ac.jp/crid/1130282269526926336

Schlögl, A., Keinrath, C., Zimmermann, D., Scherer, R., Leeb, R., & Pfurtscheller, G. (2007). A fully automated correction method of EOG artifacts in EEG recordings. Clinical Neurophysiology, 118(1), 98–104. 10.1016/j.clinph.2006.09.003

Schubring, D., & Schupp, H. T. (2019). Affective picture processing: Alpha- and lower beta-band desynchronization reflects emotional arousal. Psychophysiology, 56(8), e13386. 10.1111/psyp.13386

Schupp, H. T., Stockburger, J., Codispoti, M., Junghöfer, M., Weike, A. I., & Hamm, A. O. (2006). Stimulus novelty and emotion perception: The near absence of habituation in the visual cortex. NeuroReport, 17(4), 365. 10.1097/01.wnr.0000203355.88061.c6

