## Supplemental Figure 1 for "Emotional Responses to Naturalistic and AI-generated Affective Pictures: A Systematic Comparison"

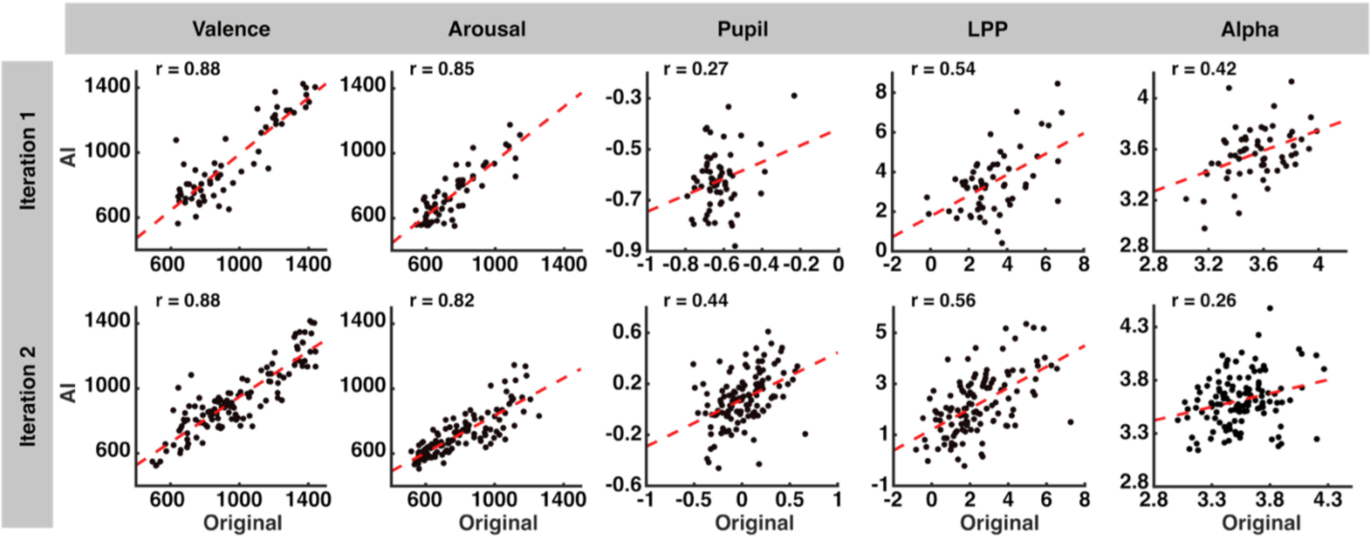


**Supplemental Figure 1.** Correlation between matched AI and original picture pairings, for both iteration 1 and iteration 2. Each dot represents a picture, and each line represents the line of best fit for the data, using a least-squares regression.
