## Supplemental Figure 2 for "Emotional Responses to Naturalistic and AI-generated Affective Pictures: A Systematic Comparison"

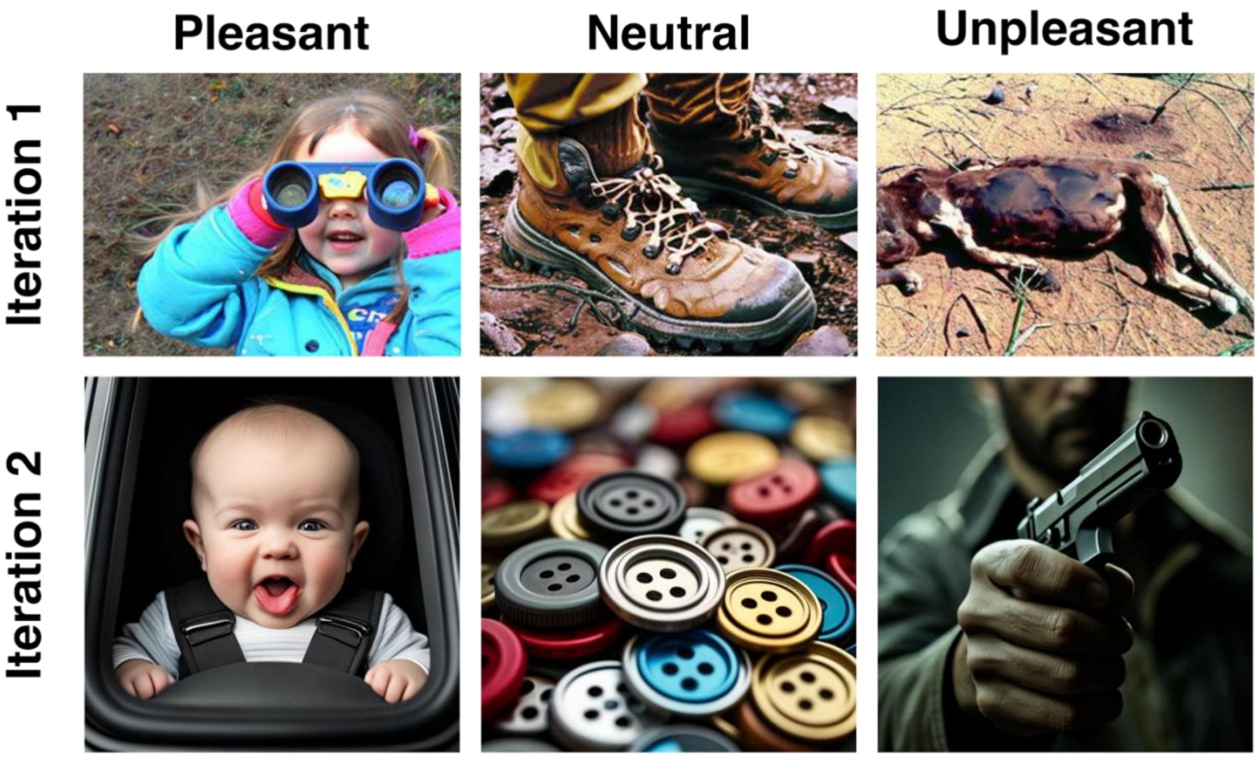


**Supplemental Figure 2.** Exemplars of AI-generated pictures for each iteration and each picture category. Example pictures are taken from the AI-generated stimuli, and are not original pictures nor real individuals.
